# Increased striatal coupling one week after ischemic stroke revealed by ultrafast functional MRI

**DOI:** 10.1101/2025.11.12.688149

**Authors:** Rita Alves, Joana Cabral, Tânia Carvalho, Noam Shemesh

## Abstract

**Background:** Network reorganization following ischemic stroke is thought to play a role in recovery. Although cortico-cortical reorganization is widely established, changes in interhemispheric striatal connections following ischemia remain poorly understood, even when stroke occurs in motor areas. Given the importance of the striatum to motor function, we investigated network-level striatal coupling in stroke using ultrafast resting-state fMRI, which has recently been shown to facilitate the dissection of synchronous oscillatory activity better than its conventional ∼1 sec time-resolution counterparts.

**Methods:** A cohort of (N=18) sedated rats were randomized and N=9 rats underwent unilateral photothrombotic ischemic lesioning in motor cortex. One week after the lesion, when plasticity and recovery are well established, all animals were scanned on a 9.4T MRI scanner using a cryogenic coil using an ultrafast resting-state functional MRI sequence with temporal resolution of 90 ms. Data were collected for 24 minutes, and spectral power, phase locking, and functional connectivity were quantified. Histology was performed to confirm lesion extent.

**Results:** While cortico-cortical power, connectivity and synchrony were diminished one week post-stroke as expected, we surprisingly found increased striato-striatal power, synchrony and functional connectivity in the stroked group compared with the control group. In stroked animals, the spectral power in the ultraslow oscillation frequency band (0.02-0.4 Hz) significantly increased in the striatum while decreasing in the cortex. When data were undersampled to “conventional” fMRI temporal resolution (900 ms), the striatal effects were lost, revealing the power of ultrafast fMRI approaches in unveiling such phenomena.

**Conclusions:** Increased striato-striatal coupling, in the form of increased synchrony, spectral power, and functional connectivity, was revealed by ultrafast resting-state fMRI, but not conventional temporal resolution resting-state fMRI. Our findings suggest more involvement of subcortical areas in network reorganization than previously thought.

## Introduction

The brain possesses a remarkable innate capacity for spontaneous recovery^1^ in stroke^2^ as well as other diseases^3^. After the acute event, adaptive changes such as axonal sprouting^4^, dendritic remodeling^5^, and synaptogenesis^5^ typically occur on multiple spatial and temporal scales. At the functional network level, reorganization in connections and/or activity patterns can partially compensate for disrupted or lost neural activity^6^, improving brain function and potentially leading to improved behavioural outcomes, although the exact relationships still remain unknown, making outcome prediction very difficult^7,8^.

Functional magnetic resonance imaging (fMRI)^9^ has become a key tool for characterizing post-stroke network reorganization. Task or stimulus-evoked fMRI studies in both humans and rodents have revealed abnormal activation patterns in stroked subjects, such as ipsilateral cortical responses to stimulation of the impaired limb, that later diminishes as contralateral function is recovered to some extent^10–12^. Resting-state fMRI (rsfMRI), which is performed without external stimulation and thereby reflects spontaneous brain activity, provides rich information on global brain networks, connectivity between areas, and insights into how they change in health and disease^13–15^. Notably, following stroke, rsfMRI studies have documented disruptions in long range correlations, or functional connectivity (FC), mainly reporting reduced interhemispheric cortical connectivity that correlate with poor motor outcomes^14,16^.

In contrast to the extensive attention paid to cortical networks, the role of subcortical structures – particularly the striatum – in stroke recovery has received much less focus, even when stroke involves motor areas. As a major component of the basal ganglia, the striatum plays a central role in motor (as well as emotional and cognitive) processing^17^. Tractography studies suggest that in large strokes transcallosal fibers link the contralateral striatum to the ischemic territory, as well as new connections between perilesional cortex and the thalamus^18^. Still, the contribution of striatal circuits to functional network reorganization remains poorly understood, in part because conventional rsfMRI analyses have struggled to robustly capture changes in subcortical dynamics.

The constraints of widely used FC metrics and the dynamic nature of resting-state activity itself^19^ motivated alternative analysis frameworks, such as co-activation patterns (CAPs)^20,21^ and quasi-periodic patterns (QPPs)^22^, that have revealed transient and recurring patterns of spontaneous brain activity. More recently, ultrafast fMRI^23,24^ – which was previously shown to resolve the order of information flow in task-induced activation across the brain – has revealed the dynamics of spontaneously recurring patterns; in particular, the high temporal resolution has allowed the detection of oscillatory activity extending throughout the cortex and the striatum, driving long-range correlations across distant parts of the brain^25^. Studies combining fMRI with optical calcium imaging pointed to direct links with neural activity^26,27^, suggesting a potential role for ultrafast fMRI in characterizing spatio-spectral properties in health and in disease.

Given the importance of the striatum in motor function^28^ we hypothesized that striatal networks would participate in post-stroke functional reorganization, particularly when ischemic lesions affect motor regions. To investigate ultraslow oscillatory properties, we harnessed in-vivo ultrafast fMRI^25^ in a photothrombotic stroke model^29^ targeting the primary motor cortex one week post-ischemia, a time point in which both extensive plasticity^30^ and pronounced behavioural improvement in motor function^10^ are well established. We found enhanced striato-striatal interactions that were undetectable using conventional temporal resolution, revealing new aspects of post-stroke network reorganization.

## Methods

### Ethical statement

All animal experiments complied with the European Directive 2010/63 (established by Portuguese Decree-Law 113/2013) and followed the Federation of European Laboratory Animal Science Associations (FELASA) guidelines and recommendations concerning laboratory animal welfare. Experiments were preapproved by the Champalimaud Foundation’s Internal Review Board (ORBEA) and by the Portuguese competent authority for animal welfare (DGAV, Direção Geral de Alimentação e Veterinária) with license number 0421/000/000/2016.

### Experimental design

In this study, N = 18 adult Long-Evans female rats weighing 311.0 ± 54.9 g were used. Animals were reared in a temperature-controlled room with *ad libitum* access to food and water and under a normal 12 h/12 h light/dark cycle. N = 9 animals were randomly selected as controls (healthy) and N = 9 were randomly assigned as the stroke group.

The experimental workflow is summarized in Figure 1. A unilateral photothrombotic cortical stroke was induced in the stroke group rats, followed by the fMRI sessions 1-week post-ischemia (Figure 1A-D).

**Figure 1.**
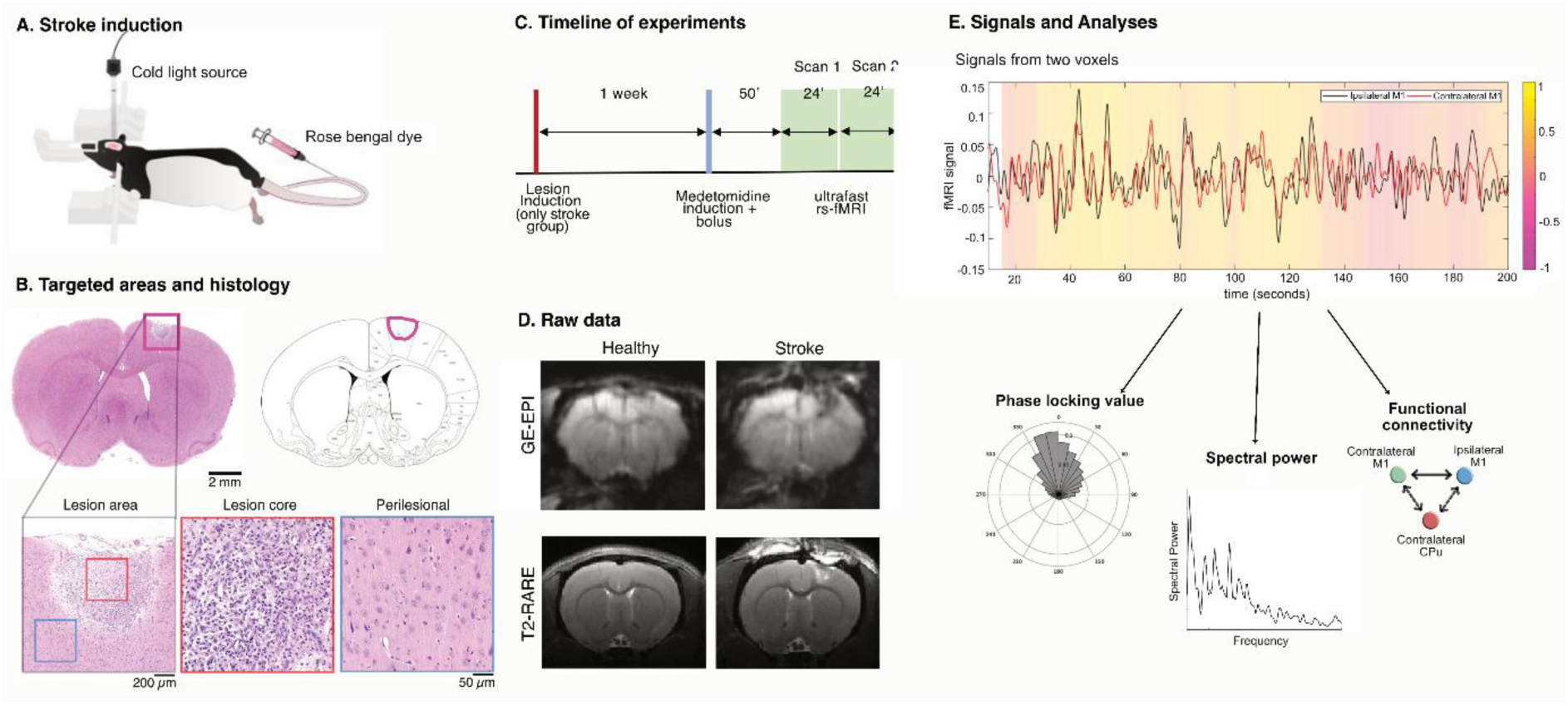
Experimental design, histology, raw data, and analysis. **(A)** A unilateral cortical stroke was performed using a photothrombotic stroke model^29^ in N = 9 rats, while 9 other rats served as controls. **(B)** Histopathological features in the targeted areas following photothrombotic stroke. A focal, well-demarcated lesion is observed in the caudal forelimb area of the primary motor cortex (cM1), at the level of the striatum (left, coronal H&E-stained section; middle, brain atlas reference). High-magnification views show the lesion core (red box) with neuronal loss, vacuolation, gliosis, and inflammatory cell infiltration, consistent with ischemic necrosis. The perilesional cortex (blue box) displays no signs of ischemic damage. Original magnification: 5x, scale bar = 200 µm (left photo); 20x, scale bar = 50 µm (red and blue photo boxes). **(C)** 1 week following ischemia, the animals were subject to MRI scans (9.4T), under medetomidine, and two ultrafast fMRI scans were performed 50 minutes after sedation induction to ensure stable sedation. **(D)** Raw data from the ultrafast fMRI scans (top row) and anatomical images (bottom row) from a representative animal reveal high quality data (left column, control; right column, stroke). **(E)** Representative time course in the ipsilateral and contralateral primary motor cortices (iM1 and cM1) from a healthy control ultrafast fMRI. A synchrony analysis was performed by calculating the phase locking value in different ROI pairs between groups; a spectral power analysis was performed to calculate the total power across different regions between groups, as well as the power across different frequency bins; a conventional functional connectivity (FC) analysis using seed-based correlation was performed and compared between groups.

### Surgical procedures

A photothrombotic Rose Bengal stroke model^29^ was used to induce a unilateral focal infarction in the motor cortex (M1) of N = 9 rats (Figure 1A). Each rat was injected with meloxicam 30 min prior to surgery and anesthetized with isoflurane (∼2.5% in oxygen-enriched medical air composed of 70% nitrogen, 29% oxygen and the remaining 1% comprising mostly argon, carbon dioxide and helium). Temperature was monitored using a rectal probe and maintained at 36 – 37 °C with a heating pad. The skull was exposed by a median incision of the skin at the dorsal aspect of the head and the skull was thinned over the right M1 (1.67 mm posterior; 2.5 mm lateral to bregma^31^. A solution of Rose Bengal dye (Sigma Aldrich, 95%) dissolved in sterile saline at a 15 mg/ml concentration and filtrated through a 0.2 μm sterile filter was delivered intravenously by the tail vein (2 ml/kg body weight). The brain was then irradiated with a cold light source through a fiber optic light guide (0.89 mm tip) (8065812001, Alcon, USA) - reaching a color temperature of 3200 K and a beam light intensity of 10 W/cm^2^ – for 15 min. After the surgery, the incision was sutured and the animal was injected with 5 mg/Kg body weight of carprofen (Rimadyl, Zoetis, USA) each 24h for 3 days. The animals were then returned to their cages and allowed to recover for one week.

### Animal preparation for MRI

One week following stroke, the animals were removed from their home cage and anesthesia was induced with 5% isoflurane. The rats were weighed, moved to the MRI bed and isoflurane was reduced to 2-3%. Approximately 5 minutes after induction, a bolus of medetomidine solution (1:10 dilution of 1 mg/ml medetomidine solution (Vetpharma Animal Health S.L., Barcelona, Spain) in saline) was administered by subcutaneous injection (0.05 mg/kg) via a syringe pump (GenieTouch, Kent Scientific, Torrington, Connecticut, USA), immediately followed by the initiation of a subcutaneous constant infusion of 0.1 mg/kg/h medetomidine. Isoflurane concentration was gradually decreased to 0% over the next 10 min, while the medetomidine infusion was maintained constant throughout the remainder of the MRI session. During the entire experiment, animals breathed oxygen-enriched (29%) medical air.

Temperature and respiratory rate were monitored using a respiration pillow sensor and an optic fiber rectal temperature probe (SA Instruments Inc., Stony Brook, USA). Each experiment lasted ∼2h. At the end of the experiment, the rats were removed from the scanner and a 5 mg/ml solution of atipamezole hydrochloride (Vetpharma Animal Health, S.L., Barcelona, Spain) diluted 1:10 in saline was injected subcutaneously with the same volume as for the medetomidine bolus to reverse the sedation.

### MRI scanning

MRI scans were performed using a 9.4 T BioSpec MRI scanner (Bruker Biospin, Ettlingen, Germany) equipped with an AVANCE IIIHD console, producing isotropic pulsed field gradients of up to 660 mT/m with a 120 μs rise time and operating Paravision 6.01 software (Bruker Biospin, Ettlingen, Germany). RF transmission was achieved using an 86 mm quadrature resonator, while a 4-element array cryoprobe (Bruker, Fallanden, Switzerland) was used for signal reception. Following localizer experiments and routine adjustments for center frequency, RF calibration, acquisition of B_0_ maps, and automatic shimming, anatomical images were acquired using a T_2_-weighted RARE sequence in the coronal and sagittal planes: TR/TE = 2000/36 ms, FOV = 18 × 16.1 mm^2^ (coronal) or FOV = 24 × 15.5 mm^2^ (sagittal), in-plane resolution = 150 × 150 μm^2^ (coronal) or 150 × 145 μm^2^ (sagittal), RARE factor = 8, slice thickness = 0.6 mm (coronal) or 0.5 mm (sagittal), number of slices: 22 (coronal) or 21 (sagittal), t_acq_ = 3 min 28 s. These images were used to guide the positioning of the ultrafast fMRI single-slice acquisitions.

### Ultrafast fMRI acquisitions

Ultrafast fMRI focuses on a single slice to ensure very high repetition rates^23,25^. In this study, a 1.2 mm-thick slice covering both cortical and striatal areas of the rat brain was chosen. The slice was centered at 0.8 mm from Bregma^31^. Two ultrafast fMRI scans (spontaneous activity) were acquired for each of the N = 18 rats (totaling 36 scans) using a gradient-echo echo planar imaging (GE-EPI) sequence (TR/TE = 90/16 ms, flip angle = 20°, FOV of 21 × 21 mm^2^, matrix size = 84 × 84, in-plane resolution = 250 × 250 μm^2^, number of time frames = 14000, t_acq_ = 24 min, and dummy scans = 2023 to ensure that steady magnetization and amplifier stability are achieved.

### Data analysis

Data analysis was performed using custom written code in MATLAB (The Mathworks Inc., Natick, MA, USA, v2018a and v2021a), ITK-Snap^32^ and ImageJ^33^. Group comparisons used two-sided t-tests with FDR control (Benjamini–Hochberg, q=0.05) for multiple comparisons.

### Brain masking and animal alignment

Individual brain masks were manually drawn and aligned across rats to a common central coordinate. All individual rat masks were superposed to define a common brain mask containing N_v_ = 1263 voxels in the single fMRI slice.

All datasets underwent realignment via a sub-pixel registration method^34^, in which each frame was aligned to a common reference frame (first frame) from the same acquisition.

### Data denoising

The fMRI signals in the N_v_ = 1263 coronal brain voxels were detrended, a manual outlier correction was performed (frames with signal intensity 3 times higher or lower than the standard deviation of the entire time course were replaced using spline interpolation taking the entire time course, <1% of data were identified as outliers,). The signals were then band-pass filtered between 0.02-0.4 Hz and a Principal Component Analysis (PCA) of the data was performed^25^: for each individual fMRI scan, the N_v_×N_v_ covariance matrix was computed and then the covariance matrices were averaged across the 18 scans in each group. The optimal number of components was determined based on the elbow method^35^. The first 10 eigenvectors were extracted and mapped (Figures S1 and S2), and the data was effectively denoised by obtaining a low-rank projection using only these first 10 principal components. These denoised data signals were then used downstream for all analyses described below.

### Space-frequency analysis of fMRI data

Power spectra were computed voxelwise from the denoised fMRI signals using the fast Fourier transform (FFT). To account for inter-subject variability, total power values were z-score normalized across voxels. A total power map in the filtered frequency range (0.02 – 0.4 Hz) was calculated by averaging the normalized broadband power in each voxel across all 18 scans. Images of the power across a selected range of frequencies were obtained by averaging power in specific bins in each voxel across all scans. For regional quantification, voxel-wise values within each anatomically defined ROI were averaged to yield a single mean z-scored total power value per scan and per ROI. All spectral analyses were performed at the single scan level and results were then averaged for each group (N=18 spectra per group).

### Synchrony analysis of fMRI data

To assess phase synchrony between brain regions, the denoised time series were first transformed into the analytical signal using a Hilbert transform^36^, from which the instantaneous phase was extracted. For each scan, the phase time series were mapped voxel-wise, and then spatially averaged within manually defined ROIs (based on the SIGMA Atlas template^37^: contralateral motor cortex (cM1), ipsilateral primary motor cortex (iM1), contralateral primary forelimb somatosensory cortex (cS1FL) and ipsilateral primary forelimb somatosensory cortex (iS1FL), contralateral caudate putamen (cCPu), and ipsilateral caudate putamen (iCPu)) by computing the circular mean of the complex-valued phase signal. Phase synchrony between ROI pairs (cM1 and iM1, cM1 and cS1FL, cM1 and iS1FL, cM1 and cCPu, cM1 and iCPu, iM1 and cS1FL, iM1 and iS1FL, iM1 and cCPu, iM1 and iCPu, cS1FL and iS1FL, cS1FL and cCPu, cS1FL and iCPu, iS1FL and cCPu, iS1FL and iCPu, and cCPu and iCPu) was then quantified using the Phase Locking Value (PLV)^38^, calculated as the magnitude of the mean of the complex phase difference vectors across time:

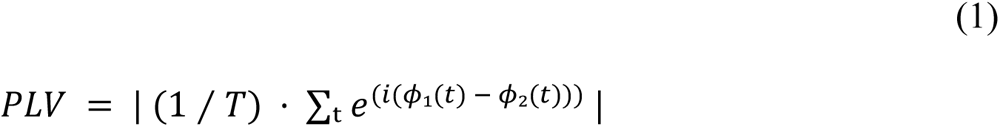

where *T* is the number of samples, and 𝜙_*i*_ is the signal phase of component ‘*i*’ derived from the Hilbert transform. PLV was calculated separately for each scan and then averaged per group.

### FC Analysis

Resting-state fMRI denoised signals within the brain mask were used for FC analyses. ROI seed-based (Pearson’s) correlation maps were generated from filtered data using cM1, iM1, cS1FL, iS1FL, cCPu and iCPu. Correlation coefficients were converted to Fisher r-to-z values before group-level comparisons. Group maps were computed from the averaged seed-based correlation maps averaged between a total of N = 18 scans per group.

### Downsampling in time to more conventional rsfMRI temporal resolution

When indicated, data were subsequently subsampled to a temporal resolution of 900 ms to compare the results from our analyses between ultrafast acquisitions (TR = 90 ms) and a lower time resolution dataset representing more conventional rsfMRI scans.

### Histopathology validation

In N=2 rats, whole brains were collected after transcardial perfusion-fixation with 4% paraformaldehyde, paraffin-embedded, and sectioned at 4 μm thickness. Coronal sections spanning the area of the ischemic lesion were stained with hematoxylin and eosin (H & E; Sigma-Aldrich, St. Louis, MO, USA) and examined by a board-certified veterinary pathologist in a Zeiss Axioscope 5 microscope equipped with an Axiocam 208 camera. Extent of the lesion and associated morphological changes were assessed in representative regions of the affected tissue.

## Results

### Lesion induction and data quality

Pathological analysis 1 week post ischemic onset revealed a focal, well-demarcated lesion in the forelimb area of the primary motor cortex (cM1) (Figure 1B). On high-magnification (red box, Figure 1B), the lesion core was characterized by extensive neuronal loss, vacuolation, gliosis, and infiltration of inflammatory cells, indicative of ischemic necrosis. Perilesional cortex (blue box, Figure 1B) is shown for comparison, depicting preserved morphological features with no significant lesions.

Figure 1D shows raw data from ultrafast fMRI in a representative animal from the healthy group (left) and from the stroke group (right). Good brain contrast, few EPI distortions, and no discernible motion artifacts (c.f. Supplementary Raw_Movie1) were noted. The SNR in the intact cortical, contralesional striatal and ipsilesional ROIs were, respectively: 28.48±2.61 (healthy) and 27.34±5.56 (stroke) (*P* = 0.4723), 19.51±1.78 (healthy) and 17.54±3.41 (stroke) (*P* = 0.0725), 18.78±1.63 (healthy) and 17.86±3.97 (stroke) (*P* = 0.4088) (reported as mean ± SD) in the ultrafast fMRI datasets.

### Cortical spectral power is reduced but striatal spectral power is increased 1 w post-stroke

Spectral power was examined in multiple areas (Figure 2), both at between 0.02 – 0. 4 Hz and at a narrower band level, across 8 distinct, non-overlapping frequency bands (Figure S3). The power spectra (Figure 2A) and maps derived from ultrafast fMRI data (Figure 2B) revealed striking differences between the healthy and stroke groups.

**Figure 2.**
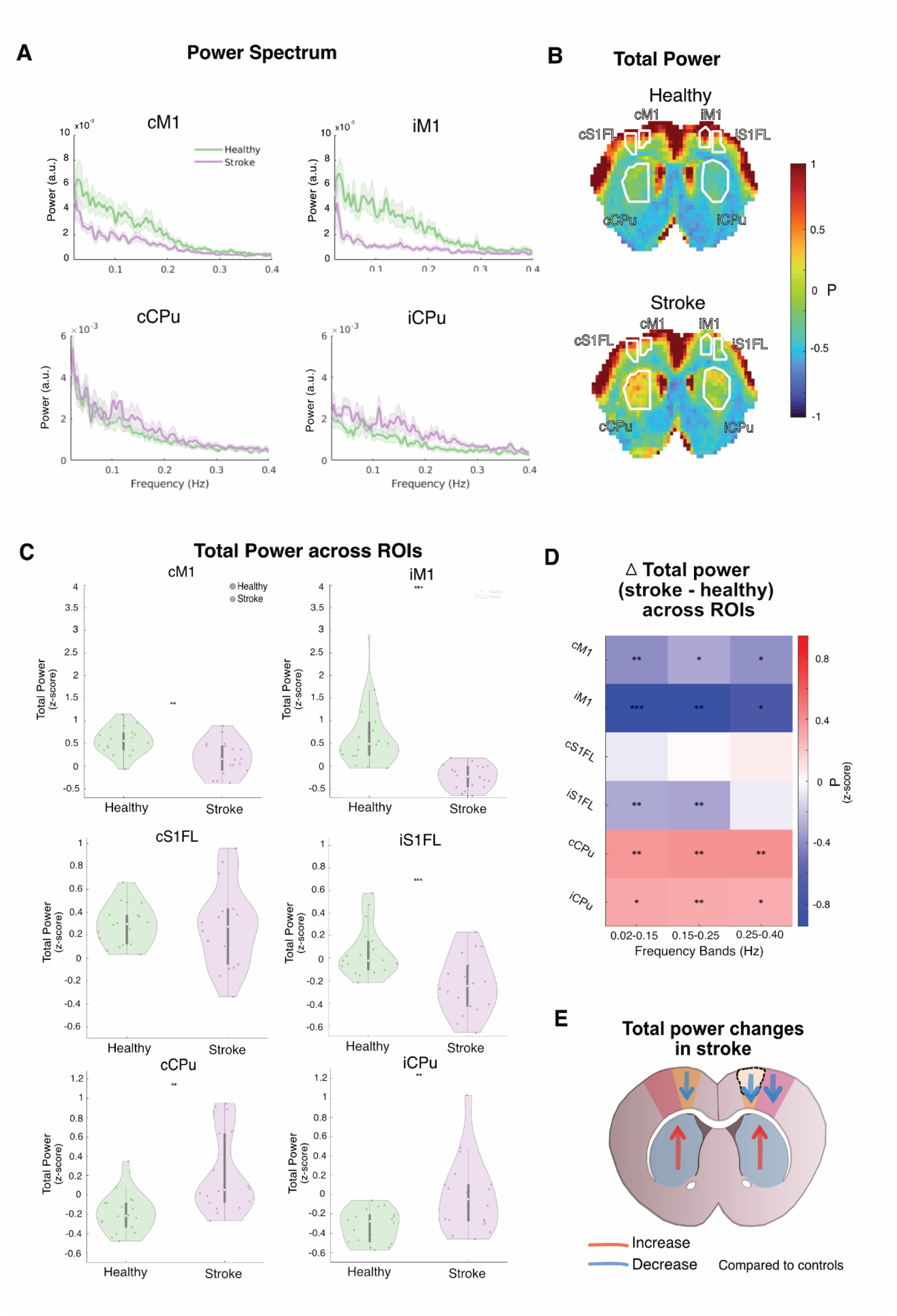
Power analysis of ultrafast fMRI signal. **(A)** Mean power spectra of the denoised ultrafast fMRI signal between 0.02 and 0.4 Hz across four ROIs: cM1, iM1, cCPu and iCPu. Shaded areas represent SEM across animals. The healthy group (green) show higher power between 0.02 and 0.4Hz compared to the stroke group (purple), particularly in cortical regions, whereas higher total power is observed at the subcortical regions in the stroke group. **(B)** Group-averaged spatial maps of total power (0.02–0.4 Hz range) in healthy and stroke rats. Cortical areas exhibit reduced power in the stroke group, while increased power is seen in subcortical areas, particularly in cCPu. **(C)** Violin plots showing z-score normalized total spectral power within each ROI: cM1, iM1, cS1FL, iS1FL, cCPu, and iCPu. Cortical ROIs exhibit significantly reduced power in stroke animals, while striatal ROIs (cCPu and iCPu) show increased power. Each green (healthy) and purple (stroke) dot represents a scan from an individual animal; white dots mark group medians. Statistical comparisons used two-sample t-tests with FDR correction (* *P* < 0.05; ** *P* < 0.01; *** *P* < 0.001). **(D)** Heatmap of ΔPower (Stroke - Healthy) across ROIs and 3 frequency bands (0.02 – 0.15 Hz; 0.15 – 0.25 Hz; 0.25 – 0.4 Hz). Warmer colors indicate increased normalized power in stroke animals, while cooler tones indicate reductions. Asterisks denote statistically significant differences after FDR correction. A reduction in normalized power is observed in iM1, particularly in between 0.02 and 0.25Hz, whereas cCPu and iCPu exhibit increased power, most prominently between 0.15 and 0.25 Hz. **(E)** Schematic summary of stroke-related changes in power. Red arrows denote regions with increased power in stroke, and blue arrows indicate regions with reduced power compared to healthy controls.

Notably, power spectra revealed decreased total power across ipsilateral and contralateral cortical motor regions, while conversely, the ispsilateral and contralateral striata clearly exhibits increased power. The maps also clearly show these trends, and they are further confirmed in the ROI-based quantification (Figure 2C), where the broadband power was significantly reduced in M1 on the lesion side following stroke, cM1 (*P* = 0.0029) and iM1 (*P* = 0.0002), and significantly increased in cCPu (*P* = 0.0017) and iCPu (*P* = 0.0041). Figure 2D parses power into 3 bins for multiple ROIs, confirming that cortical areas exhibit decreased power (blue) and striatal areas increased power (red) in most spectral areas between 0.02-0.4Hz, with the exception of cS1FL and iS1FL at the higher power bin. Figure 2E schematically represents these findings with blue arrows reflecting decreased power and red arrows increased power.

### Cortico-cortical synchrony is reduced but striato-striatal synchrony is higher 1w post ischemia

To interpret these power changes, we next examined phase synchrony by analyzing the PLV across different ROI pairs. Figure 3A shows PLV maps for 3 different seeds (all contralateral), clearly showing decreased PLV in cortex but increased PLV in striatum. The corresponding polar plots of the phase clearly show loss of synchrony in the ipsilateral cortices but it is more difficult in these plots to discern the striatal effects (Figure 3B). When synchrony was quantified ROI-wise (Figure 3C), statistically significant interhemispheric PLV differences were found both in interhemispheric cortical ROIs cM1–iM1 (*P* = 1 × 10^−6^), cM1–iS1FL (*P* = 1 × 10^−11^), iM1–cS1FL (*P* = 0.0023) and cS1FL–iS1FL (*P* = 2 × 10^−6^) and in intra-hemispheric cortical connections, such as iM1 – iS1FL (*P* = 3 × 10^−7^).

**Figure 3.**
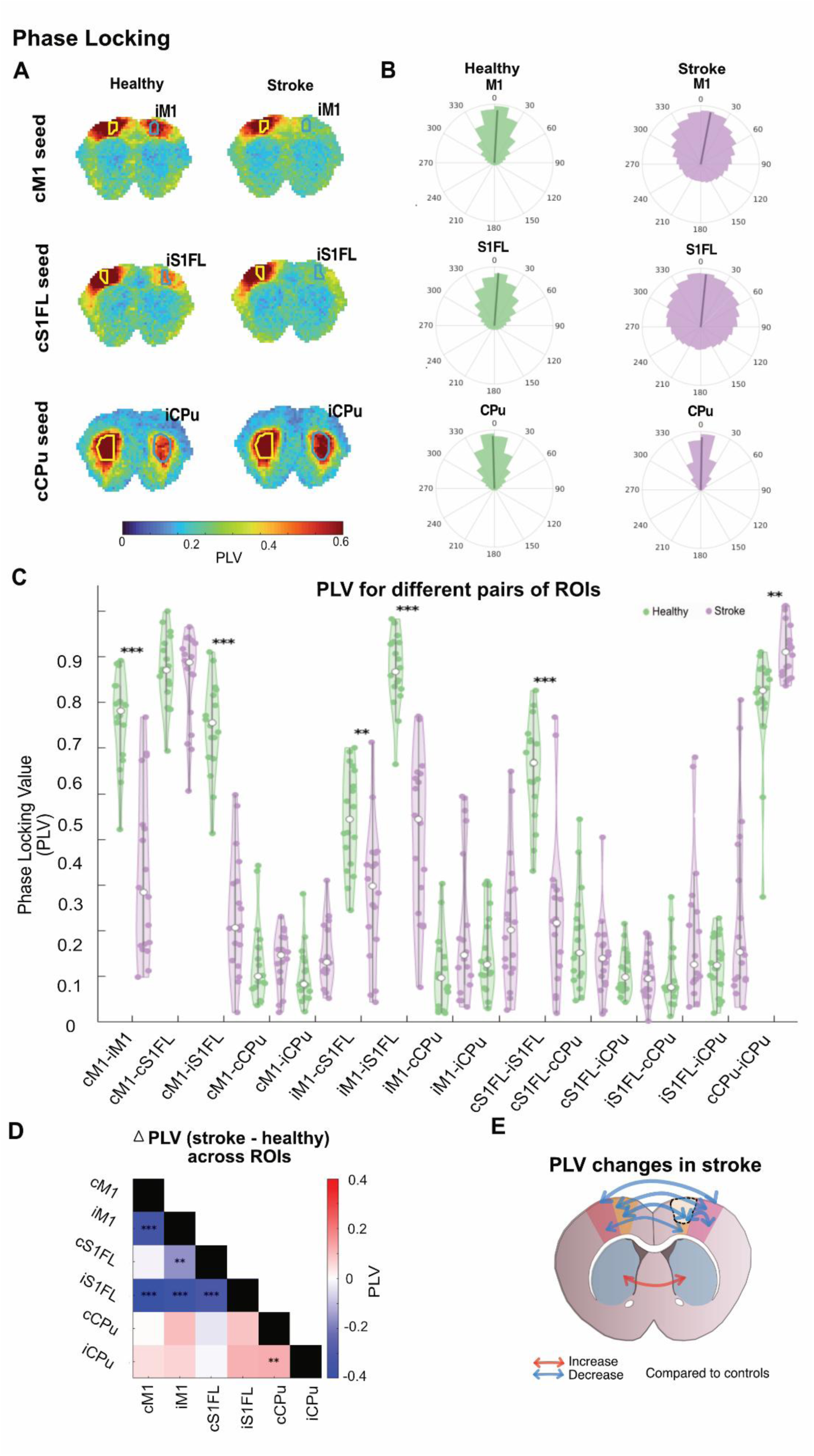
Reduced synchrony between interhemispheric cortices and increased synchrony between striatum areas 1 week after stroke. **(A)** Group-averaged Phase Locking Value (PLV) maps across selected seed regions in healthy (left) and stroke (right) animals, including different ROI pairs (cM1 and iM1, cM1 and cF1SL, iM1 and iS1FL, iM1 and cCPu, iM1 and cCPu, cM1 and cS1FL, cM1 and iS1FL, cM1 and cCPu cM1 and iCPu, cS1FL and iS1FL, cS1FL and cCPu, cS1FL and iCPu, iS1FL and cCPu, iS1FL and iCPu, and cCPu and iCPu). Warmer colors indicate higher PLV values, reflecting greater synchrony. **(B)** Polar plots representing the angular distribution of phase relationships for each ROI, showing tighter phase clustering in the stroke group for striatal ROIs and in the healthy group for cortical seeds. **(C)** Statistical analysis of PLV for different pairs of brain regions. Cortico-cortical synchrony is generally higher in healthy rats, while striato-striatal synchrony is elevated in stroke rats. White dots indicate group medians; Green and purple dots represent individual scans. Statistical comparisons use two-sample t-test with FDR correction (**\****P* < 0.05, **\*\****P* < 0.01, **\*\*\****P* < 0.001). **(D)** Heatmap of ΔPLV (Stroke – Healthy) across all ROI pairs. Warmer colors indicate increased synchrony between ROI pairs in stroke animals, while cooler tones indicate reductions. Asterisks denote statistically significant differences. **(E)** Schematic summarizing key changes in PLV in the stroke group compared to controls. Red arrows indicate increased synchrony (notably within and across striata), and blue arrows indicate decreased synchrony (primarily between cortical regions).

By contrast to this cortico-cortical uncoupling, interhemispheric coupling between the two striatum areas increased in a statistically significant fashion. Notably, cCPu – iCPu PLV was higher in the stroked group compared with controls (*P* = 0.0033, Figure 3C). Figure 3D quantifies the difference in PLV between groups for the different ROIs further highlighting the decreased cortical PLV (blue) and increased striatal PLV (red), and Figure 3E schematically represents these effects with blue arrows reflecting decreases in PLV and red arrows reflecting increases. A complementary phase-occupancy analysis (Figure S4) yielded a similar pattern, with cortical ROI pairs spending less time and striatal ROI pairs more time within ±π/6 in stroke animals, consistent with the PLV findings.

### Ultrafast fMRI reveals decreases in cortico-cortical but increases in striato-striatal functional connectivity

Next, we investigated the impact of these effects on FC (Figure 4). Figure 4A shows maps obtained from two cortical seeds and one striatal seed. In the healthy group, when cortical seed regions (cM1, cS1FL) were selected, we found strong interhemispheric connectivity (∼0.78 between cM1 and iM1; ∼0.77 between cM1 and iS1FL, ∼0.57 between cS1FL and iM1; ∼0.68 between cS1FL and iS1FL). In the stroke group, however, reduced interhemispheric connectivity was observed (∼0.33 between cM1 and iM1 (*P*= 9 × 10^−8^); ∼0.24 between cM1 and iS1FL (*P* = 6 × 10^−11^); ∼0.3 between cS1FL and iM1 (*P* = 0.0002); ∼0.29 between cS1FL and iS1FL (*P* = 9 × 10^−7^)).

**Figure 4.**
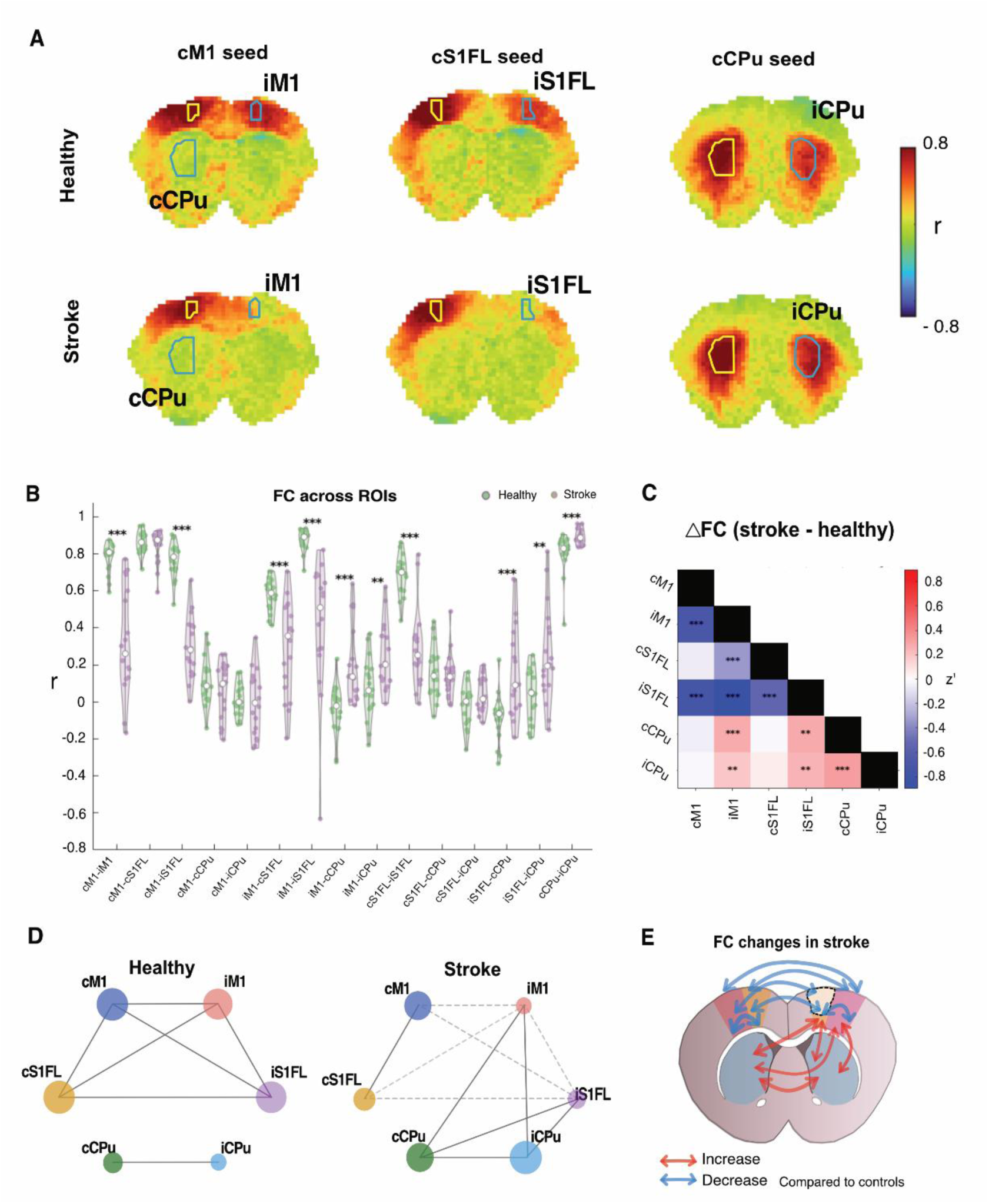
Alterations in ‘static’ resting-state functional connectivity 1 week post stroke. **(A)** Analysis in band-pass filtered fMRI signals (0.02 – 0.4 Hz) in healthy and stroke groups. Seed-based correlation maps are represented for four different seeds (yellow ROIs), where each voxel is colored according to its degree of correlation with the seed. The ROIs used to correlate with each seed are represented in blue. Correlation values were truncated between −0.8 and 0.8 for display. **(B)** Violin plots of Pearson correlation values across all ROI pairs for both groups. Cortico-cortical FC is significantly reduced in stroke rats, while striato-striatal and cortico-striatal FC is increased. Each green (healthy) and purple (stroke) dot represents a scan from an individual animal. Median values are shown as white dots. Group differences assessed using two-sample t-tests with Benjamini-Hochberg FDR correction applied to Fisher’s r-to-z values. (* *P* < 0.05; ** *P* < 0.01; *** *P* < 0.001). **(C)** Heatmap showing differences in FC (stroke - healthy) across all ROI pairs. Warmer colors in subcortical ROIs indicate increased connectivity in stroke rats; cooler colors in cortical ROIs represent decreases. Asterisks denote statistically significant differences (* *P* < 0.05; ** *P* < 0.01; *** *P* < 0.001; t-test with FDR correction). **(D)** Simplified network graphs showing FC strength for healthy and stroke groups, averaged across all ROI pairs. Edge width reflects the region’s weight (absolute connectivity strengths for each ROI). Dashed lines in the stroke group indicate weakened connections relative to controls. **(E)** Schematic illustrating FC changes observed in stroke compared to controls. Red arrows denote increases, especially in striato-striatal and cortico-striatal pairs; blue arrows indicate decreases, notably between homologous cortical areas.

Interestingly, when a seed was placed in the cCPu in the control group, high functional connectivity with its interhemispheric counterpart iCPu was found (∼0.8). In the stroke group however, connectivity values were significantly higher values between both striatum areas (∼0.89, *P* = 0.0009).

Figure 4B quantifies these effects, and Figure 4C shows the corresponding ΔFC values. We find statistically significant reductions between interhemispheric cortices and increases in the striato-striatal connectivity, and interestingly also increases in striatal connectivity to the iS1FL. Figures 4D and 4E summarize these effects schematically, showing the changes in connectivity 1w post-ischemia.

### Resting-state fMRI with conventional temporal resolution fails to detect most striatal effects

Figure 5 summarizes results for spectral power, synchrony, and seed-based FC analyses from data that was subsampled to represent sampling in conventional rsfMRI experiments (namely the effective TR was now 900 ms).

**Figure 5.**
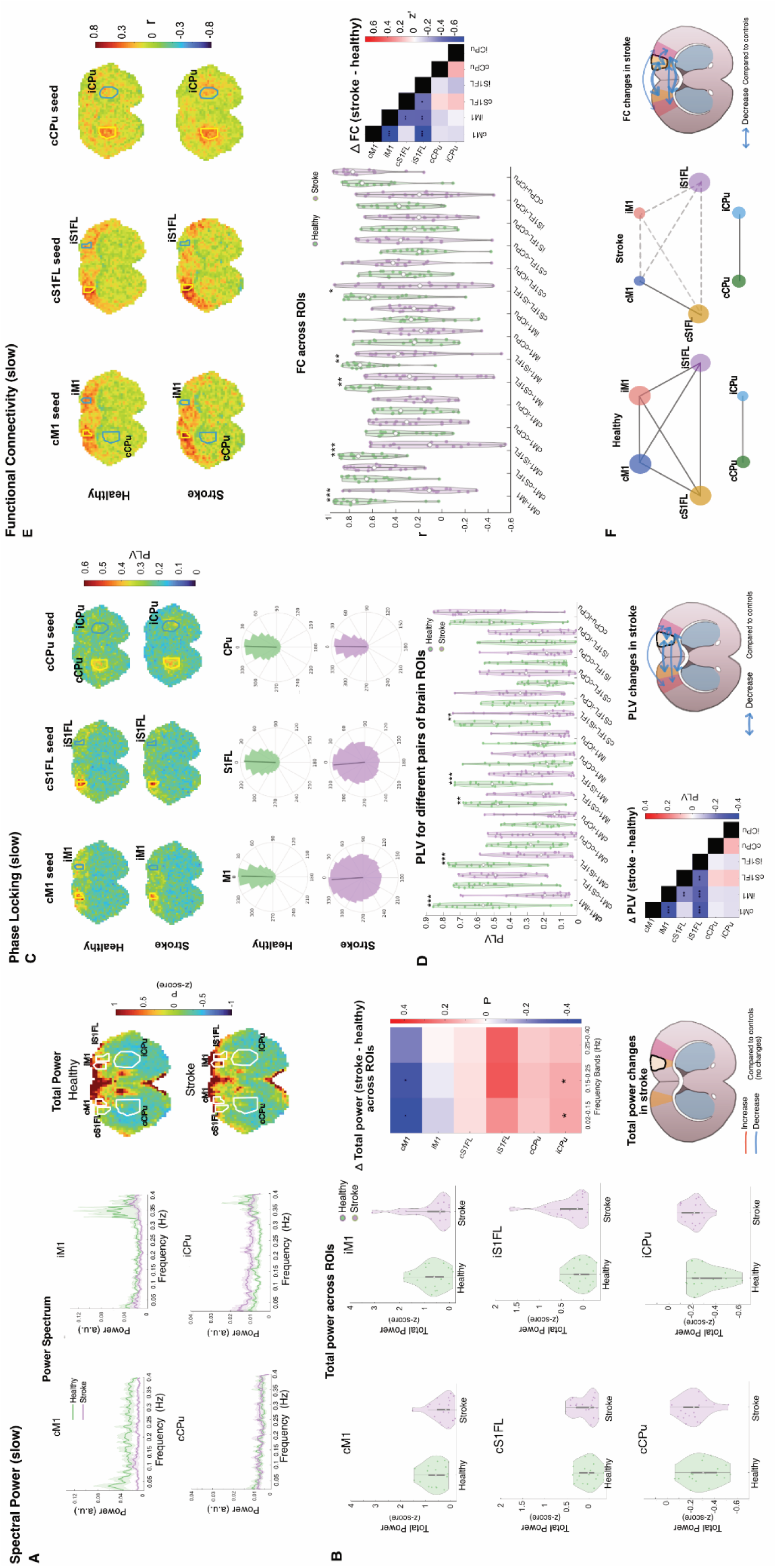
Resting-state power, synchrony, and functional connectivity analysis in conventional acquisitions (TR = 900 ms). (A, B) Power spectral analysis. **(A)** Mean power spectra for cortical ROIs show globally reduced power in stroke. Maps show spatial distribution of total power (0.02 - 0.4 Hz), with no evident alterations in stroke. **(B)** Violin plots of z-score normalized total power per ROI. Increased iCPu power was observed. Heatmap of ΔPower (Stroke – Healthy) across ROIs and frequency bands show increased iCPu power in all frequency bands. **(C, D) PLV analysis. (C)** Group-averaged PLV maps for selected seeds show reduced synchrony in stroke animals. Polar plots of phase distributions show more dispersed phase relationships in stroke. **(D)** Violin plots of PLV per ROI pair; significant reductions observed in stroke rats. Heatmap of PLV differences (stroke – healthy) across ROI pairs and schematic summary of synchrony changes show cortical decrease in PLV. **(E, F) Seed-based functional connectivity. (E)** Correlation maps from representative seeds illustrate reduced connectivity in stroke rats. Violin plots of ROI-pairwise FC and the heatmap of FC differences (stroke - healthy) exhibit a significant reduction in FC in the stroke group. **(F)** Network graphs show weakened cortico-cortical FC in stroke. Schematic summarizing FC alterations show cortical reductions in stroke. A two-sample t-test was performed for all analyses and FDR correction was used to control for multiple comparisons (\**P* < 0.05, ** *P* < 0.01).

In these subsampled data, spectral power analysis (Figure 5A–B) was not able to detect the decreases in cortical power in the cortex nor increases in striatal ROIs with statistical significance. Similarly, the maps did not clearly show any increased power in the 0.02-0.4 Hz band.

PLV-based synchrony maps (Figure 5C) revealed small differences between the healthy and stroke groups across all seeds. In the polar plots, phase clustering appeared less defined in both groups compared to the ultrafast results. The violin plots (Figure 5D) confirmed that although cortico-cortical synchrony was reduced in stroke, subcortical differences failed to reach statistical significance (*P* > 0.05). The PLV difference matrix showed a reduction in cortical synchrony in stroke animals (e.g., cM1–iM1: - 0.33, *P* = 0.0002; cM1 – iS1FL: - 0.3, *P* = 0.0001; iM1 – cS1FL: - 0.18, *P* = 0.0054; iM1 – iS1FL: - 0.26, *P* = 0.0002; cS1FL – iS1FL: - 0.23, *P* = 0.0054). However, synchrony increases in subcortical areas observed at TR = 90 ms were not observed (not significant, *P* >0.05) at TR = 900 ms.

Seed-based FC maps (Figure 5E) managed to detect the reduced interhemispheric cortical connectivity in stroke rats, particularly between homologous M1 and S1FL regions, albeit with weak correlation values (cM1–iM1: 0.19; cS1FL–iS1FL: 0.22). The violin plots and difference matrix (Figure 5E) confirmed significant group differences in cortical ROI pairs (cM1–iM1: *P* = 0.0002; cS1FL–iS1FL: *P =* 0.0499), but these subsampled data were not able to detect striatal and cortico-striatal differences (*P* >0.05). Network graphs and the schematic summary (Figure 5F) visually highlight the limited sensitivity of conventional temporal resolution to detect the network reorganization observed at higher temporal resolution.

## Discussion

Our study revealed increased striatal activity and coupling (alongside the expected decreased cortical coupling), as measured from spectral power, synchrony and functional connectivity, in the rat brain one week following ischemic stroke – a stage at which behavioral recovery is well established in this model^16,39^. These findings indicate that subcortical circuits play a more prominent role in post-stroke network reorganization than previously recognized. Moreover, the results demonstrate the necessity of ultrafast fMRI for uncovering such subcortical dynamics, which were not detectable at conventional temporal resolutions.

### Evidence converging toward increased striatal coupling one week post-ischemia

Across multiple independent metrics – oscillatory power, phase synchrony, and functional connectivity – our data consistently point to enhanced coupling within and between striatal regions following stroke. Spectral analysis revealed a clear increase in total power in the striatum, while cortical regions displayed the hallmark and well-established marked reductions^40^. Phase locking analyses showed that inter-striatal synchrony (and, to a lesser extent, ipsilateral cortico-striatal synchrony) was significantly elevated in the stroke group, contrasting sharply with the widespread loss of cortical coherence. Functional connectivity maps corroborated these findings, showing strengthened inter- and intra-striatal correlations and also enhanced coupling between striatum and perilesional cortex. Together, these convergent signatures suggest that one week post-stroke, information flow and oscillatory coordination become increasingly dominated by subcortical dynamics.

### Potential biological underpinnings of increased coupling

The increased striatal coupling likely reflects compensatory plasticity within subcortical circuits following cortical disruption. The elevated power in striatal regions is consistent with adaptive increases in excitability and synaptic scaling mechanisms that emerge after loss of cortical input^41^. Homeostatic plasticity may restore firing balance by enhancing postsynaptic responsiveness and synaptic strength, as shown in prior studies of post-ischemic adaptation^41^. Dopaminergic neuromodulation may further amplify these effects: dopamine release and D1-receptor–dependent facilitation of long-term potentiation in corticostriatal pathways are known to peak around one week after ischemia, coinciding with behavioral recovery^42,43^.

Our phase synchrony findings suggest that oscillatory coordination shifts away from cortical hubs toward the striatum, possibly as a compensatory mechanism for maintaining temporal precision in motor circuits. Although hyper-synchrony can be maladaptive in other contexts (e.g., Parkinsonian oscillations^44^), the recovery-associated timing of our findings suggests that the increased synchrony here may reflect functional re-engagement rather than pathological overcoupling^16^.

Finally, the functional connectivity increases here observed are more in agreement with potentiation of pre-existing subcortical and cortico-striatal connections, rather than on de novo circuit formation^45^. Tractography studies in mice 21 days after MCAO showing transcallosal fibers linking both striata and strengthened thalamo-striatal pathways support the notion that structural substrates exist to sustain this enhanced communication^18^.

### Necessity of ultrafast fMRI for detecting subcortical coupling

When the data were temporally downsampled to mimic more conventional rsfMRI temporal resolution (∼1 s TR), the striatal effects vanished. This highlights that ultrafast fMRI’s high temporal resolution is crucial for resolving the rapid, transient oscillations that underlie subcortical coupling. By reducing aliasing of higher-frequency physiological fluctuations, preserving phase relationships, and by virtue of the much higher number of samples, ultrafast fMRI captures dynamic synchrony and oscillatory modes that remain obscured or unresolved in more standard acquisitions. The ability to detect these fast mesoscale interactions not only provides a window into previously inaccessible subcortical plasticity but also underscores ultrafast fMRI’s potential for identifying biomarkers of recovery in longitudinal studies.

### Limitations

As in all studies, we identify several limitations in our work. The photothrombotic stroke model used here produces focal lesions that may not fully reflect the large variability encountered in spontaneous ischemic stroke; however, in our case the model’s high lesion reproducibility was a favourable aspect and the molecular differences setting it apart from other stroke models are less relevant for the purpose of MRI-based network characterisation^46^. Additionally, our analysis focused on a single time point, limiting our ability to assess how connectivity changes evolve from the acute to the chronic stage^16^. Another limitation is the single-slice nature of our ultrafast fMRI acquisition, which arises mainly from hardware limitations (excessive heating of the gradient amplifier). Future studies could employ longitudinal designs to capture the full trajectory of changes in synchrony, and could further optimize acquisition sequences and acceleration methods to enable rapid multi-slice or whole-brain imaging with minimal compromise on temporal resolution or hardware limits^47^.

## Conclusions

Increased striatal coupling one week post ischemic stroke were discovered by ultrafast fMRI. Our findings point to a larger reorganization of subcortical areas than previously thought, and highlighted the potential of ultrafast fMRI approaches for unravelling effects that cannot be resolved with conventional temporal resolution. This bodes well for future applications of ultrafast fMRI for understanding plasticity and to potentially providing more sensitive predictors of recovery.

## Acknowledgments

The authors acknowledge the vivarium of the Champalimaud Center for the Unknown, a facility of CONGENTO financed by Lisboa Regional Operational Programme (Lisboa 2020), project LISBOA01–0145-FEDER-022170, and also the Champalimaud Histopathology Platform. The authors also want to thank Ms. Mafalda Valente for assistance in the perfusion of the rat brains for histological experiments. This work was supported by FCT – Fundação para a Ciência e Tecnologia, https://doi.org/10.54499/2021.07300.BD and under the LARSyS funding https://doi.org/10.54499/LA/P/0083/2020.

## Extended Methods

### Spectral power analysis (band-specific)

Power spectra were computed voxel wise from the reconstructed fMRI signals using the fast Fourier transform (FFT). Band power values were z-score normalized across voxels for each scan. The 8 non-overlapping frequency bands were: 0.02 – 0.05 Hz; 0.05 – 0.1 Hz; 0.1 – 0.15 Hz; 0.15 – 0.2 Hz; 0.2 – 0.25 Hz; 0.25 – 0.3 Hz; 0.3 – 0.35 Hz; 0.35 – 0.4 Hz. To assess differences in power between groups across multiple ROIs (contralesional M1 cortex (cM1), ipsilesional M1 (iM1), contralesional primary forelimb somatosensory cortex (cS1FL), ipsilesional primary forelimb somatosensory cortex (iS1FL), contralesional caudate putamen (cCPu), ipsilesional caudate putamen (iCPu), we conducted independent two-sample t-tests for each ROI. A Benjamini-Hochberg false discovery rate (FDR) correction was used to control for multiple comparisons.

### Phase occupancy analysis

To characterize the temporal dynamics of phase relationships between brain regions, a phase occupancy analysis was performed. After calculating the instantaneous phase differences between each ROI pair from the Hilbert-transformed time series, the proportion of time these differences remained within a predefined window (±π/6 radians) was calculated for each scan. This measure reflects the temporal persistence of near-synchronous states and captures how long two regions stay aligned in phase. Group comparisons for both PLV and occupancy metrics were conducted using independent two-sample t-tests across scans, followed by FDR correction for multiple comparisons.

## Extended Results

### Power analysis

To further investigate total power changes, we decomposed the broadband power into distinct frequency bands (Figure S3). Spatial maps of the power across different bands (Figure S3A) reveal a consistent spatial difference between groups across all bands: healthy rats showed higher power in cortical areas, whereas stroke rats showed markedly higher power in subcortical regions.

The power ROI analysis across different frequency bins (Figure S3B) confirms that a cortical power reduction occurs at all bins between 0.02 – 0.4 Hz in cM1 (0.02 – 0.05 Hz (*P* = 0.0039), 0.05 – 0.1 Hz (*P* = 0.0076), 0.1 – 0.15 Hz (*P* = 0.0499), 0.15 – 0.2 Hz (*P* = 0.0266), 0.2 – 0.25 Hz (*P* = 0.0148), 0.25 – 0.3 Hz (*P* = 0.0097), 0.3 – 0.35 Hz (*P* = 0.0208), 0.35 – 0.4 Hz (*P* = 0.0261), in all bins between 0.02 and 0.35 Hz in iM1 (0.02 – 0.05 Hz (*P* = 0.0012), 0.05 – 0.1 Hz (*P* = 0.0012), 0.1 – 0.15 Hz (*P* = 0.0001), 0.15 – 0.2 Hz (*P* = 0.0013), 0.2 – 0.25 Hz (*P* = 0.0049), 0.25 – 0.3 Hz (*P* = 0.0076), 0.3 – 0.35 Hz (*P* = 0.0208), and in bins between 0.02 – 0.25 Hz in iS1FL (0.02 – 0.05 Hz (*P* = 0.0076), 0.05 – 0.1 Hz (*P* = 0.0028), 0.1 – 0.15 Hz (*P* = 0.0017), 0.15 – 0.2 Hz (*P* = 0.0012), 0.2 – 0.25 Hz (*P* = 0.0162).

Interestingly, while cortical power was reduced across all frequency bins, subcortical structures such as the striatum (cCPu, iCPu) exhibited the opposite pattern, with increased power in all frequency bins both in cCPu (0.02 – 0.05 Hz (*P* = 0.0148), 0.05 – 0.1 Hz (*P* = 0.0076), 0.1 – 0.15 Hz (*P* = 0.0039), 0.15 – 0.2 Hz (*P* = 0.0063), 0.2 – 0.25 Hz (*P* = 0.0076), 0.25 – 0.3 Hz (*P* = 0.003), 0.3 – 0.35 Hz (*P* = 0.0039), 0.35 – 0.4 Hz (*P* = 0.0039) and iCPu (0.02 – 0.05 Hz (*P* = 0.023), 0.05 – 0.1 Hz (*P* = 0.0139), 0.1 – 0.15 Hz (*P* = 0.0124), 0.15 – 0.2 Hz (*P* = 0.0047), 0.2 – 0.25 Hz (*P* = 0.0148), 0.25 – 0.3 Hz (*P* = 0.0132), 0.3 – 0.35 Hz (*P* = 0.0109), 0.35 – 0.4 Hz (*P* = 0.0263)

### Occupancy analysis

To better characterize the temporal properties of synchrony, we next quantified phase occupancy – the proportion of time a given ROI pair remained within a tight phase alignment window (±π/6 radians). The occupancy results in the ultrafast data revealed a similar trend: healthy animals exhibited greater temporal alignment across several ROI pairs (Figure S4 (top row)). For example, cM1–iM1, cM1–cS1FL, iM1–cS1FL, cS1FL–iS1FL showed significantly reduced phase occupancy in stroke animals ((*P* = 0.0017), (*P* = 2 × 10^−9^), (*P* = 6 × 10^−11^), (*P* = 9 × 10^−6^), respectively). Interestingly, some connections such as cM1–iS1FL, cM1–cCPu, iS1FL–cCPu, iS1FL–iCPu, cCPu–iCPu ((*P* = 0.0059), (*P* = 0.0258), (*P* = 0.00798), (*P* = 0.0371), (*P* = 0.0008), respectively) exhibited preserved or elevated occupancy in stroke, potentially reflecting compensatory engagement or aberrant coupling in subcortical circuits. Together, these findings highlight the value of dynamic synchrony metrics, revealing altered temporal coordination across cortical and subcortical regions following stroke.

## Extended Figures

**Figure S1.**
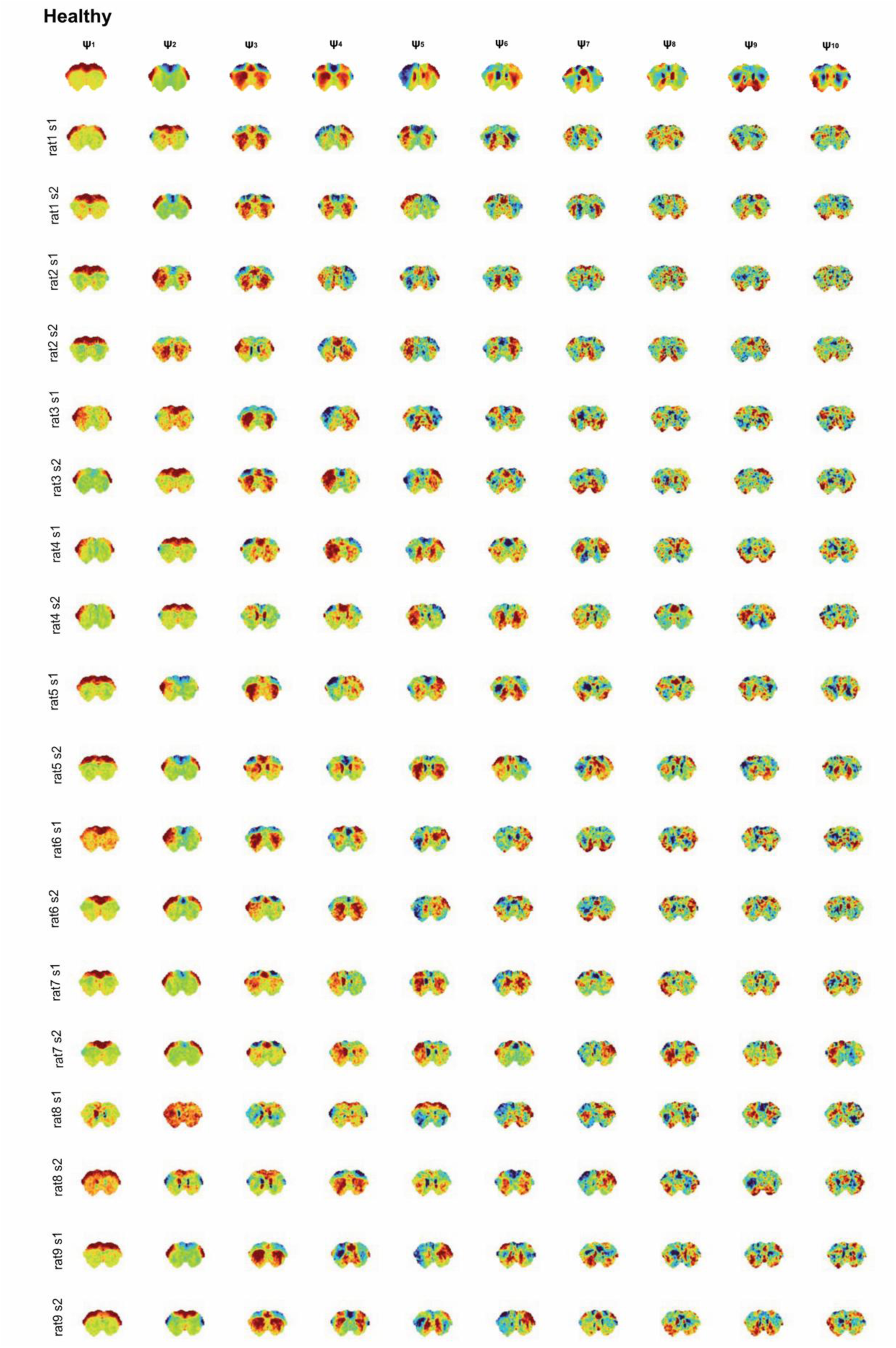
The first 10 principal components (eigenvectors) obtained from the averaged covariance matrices from the healthy group (top row), for TR = 90 ms. Striatal-striatal and cortico-striatal oscillatory patterns are observed in the stroke group in the first 2 components in the uf-fMRI dataset. Individual scans are presented below (N=18).

**Figure S2.**
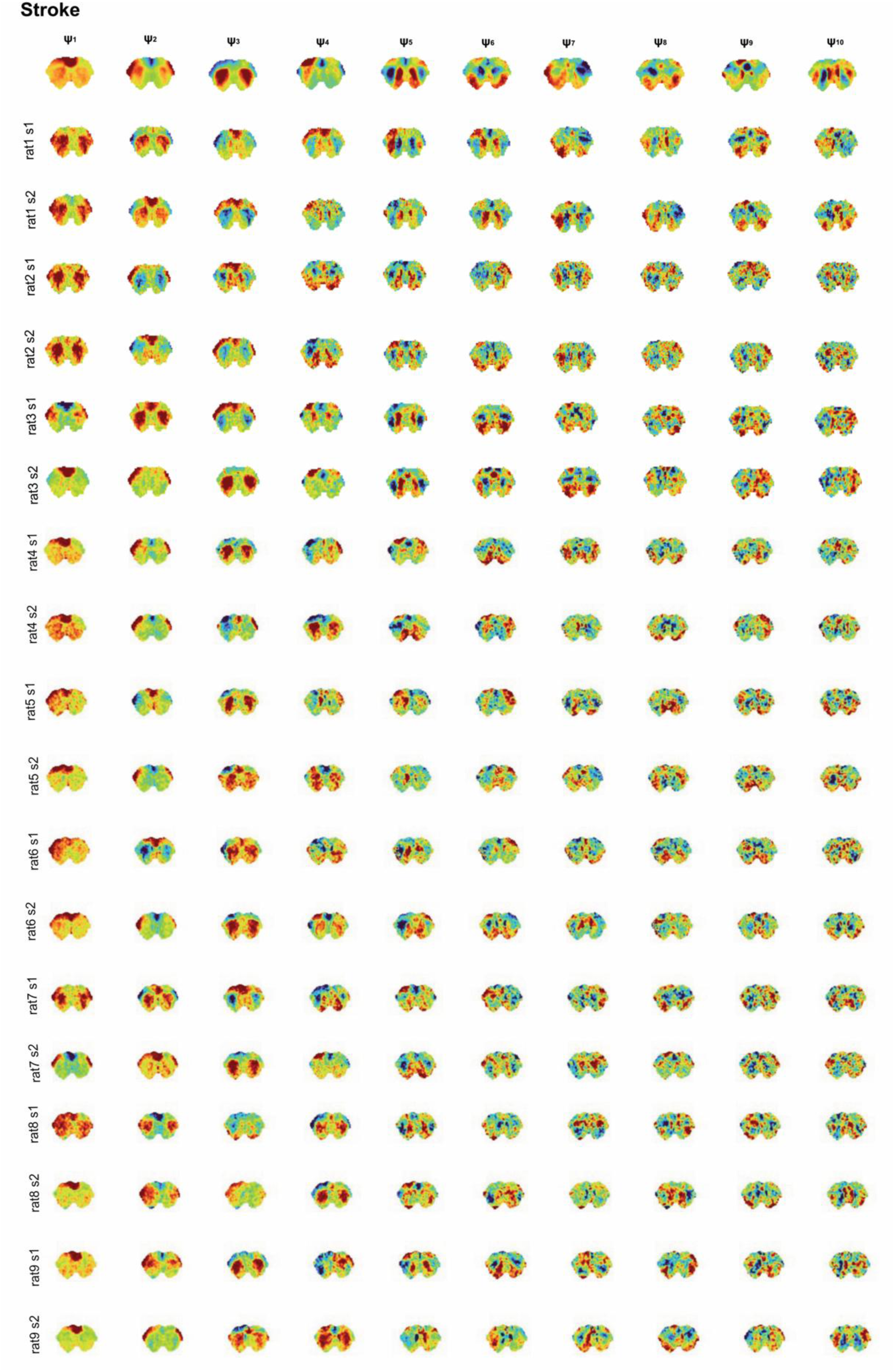
The first 10 principal components (eigenvectors) obtained from the averaged covariance matrices from the stroke group (top row), for TR = 90 ms. Striatal-striatal and cortico-striatal oscillatory patterns are observed in the stroke group in the first 2 components in the uf-fMRI dataset. Individual scans are presented below (N=18).

**Figure S3.**
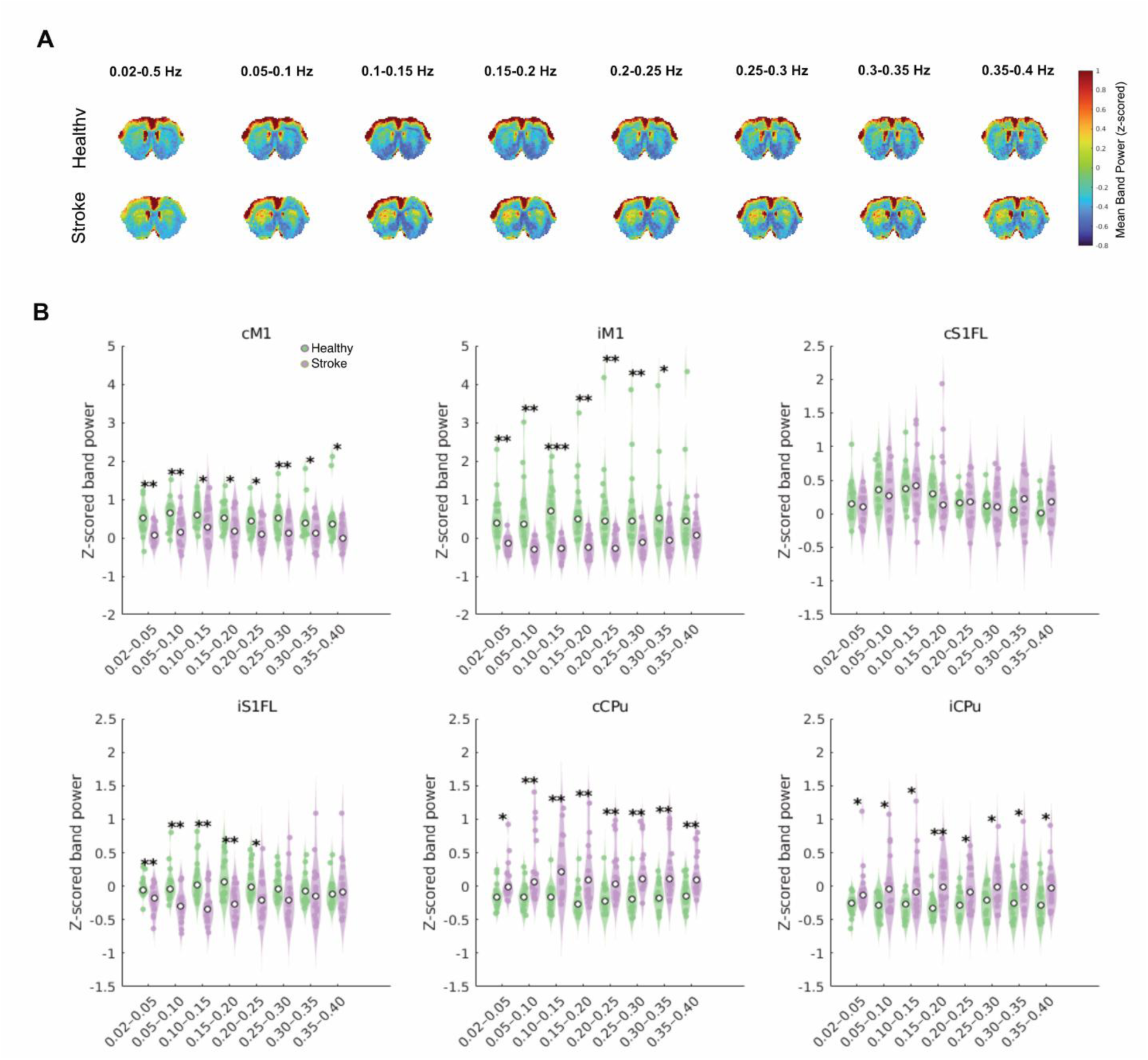
Spectral power of fMRI signal across different frequency bins. **(A)** Spatial maps of spectral power (z-score normalized) in 8 non-overlapping frequency bands, for uf-fMRI dataset, averaged across the 18 fMRI scans in each group. (B) Increased power is observed in the healthy group in comparison with the stroke group in cortical ROIs: for all frequency bins between 0.02 and 0.4 Hz in cM1, all frequency bins between 0.02 and 0.35 Hz in iM1, between 0.02 and 0.25 Hz in iS1FL. In the stroke group, higher power is observed in all frequency bins in both striata. A two-sample t-test was performed to compute differences for each ROI. A Benjamini-Hochberg FDR correction was used to control for multiple comparisons (* *P* < 0.05; ** *P* < 0.01; *** *P* < 0.001).

**Figure S4.**
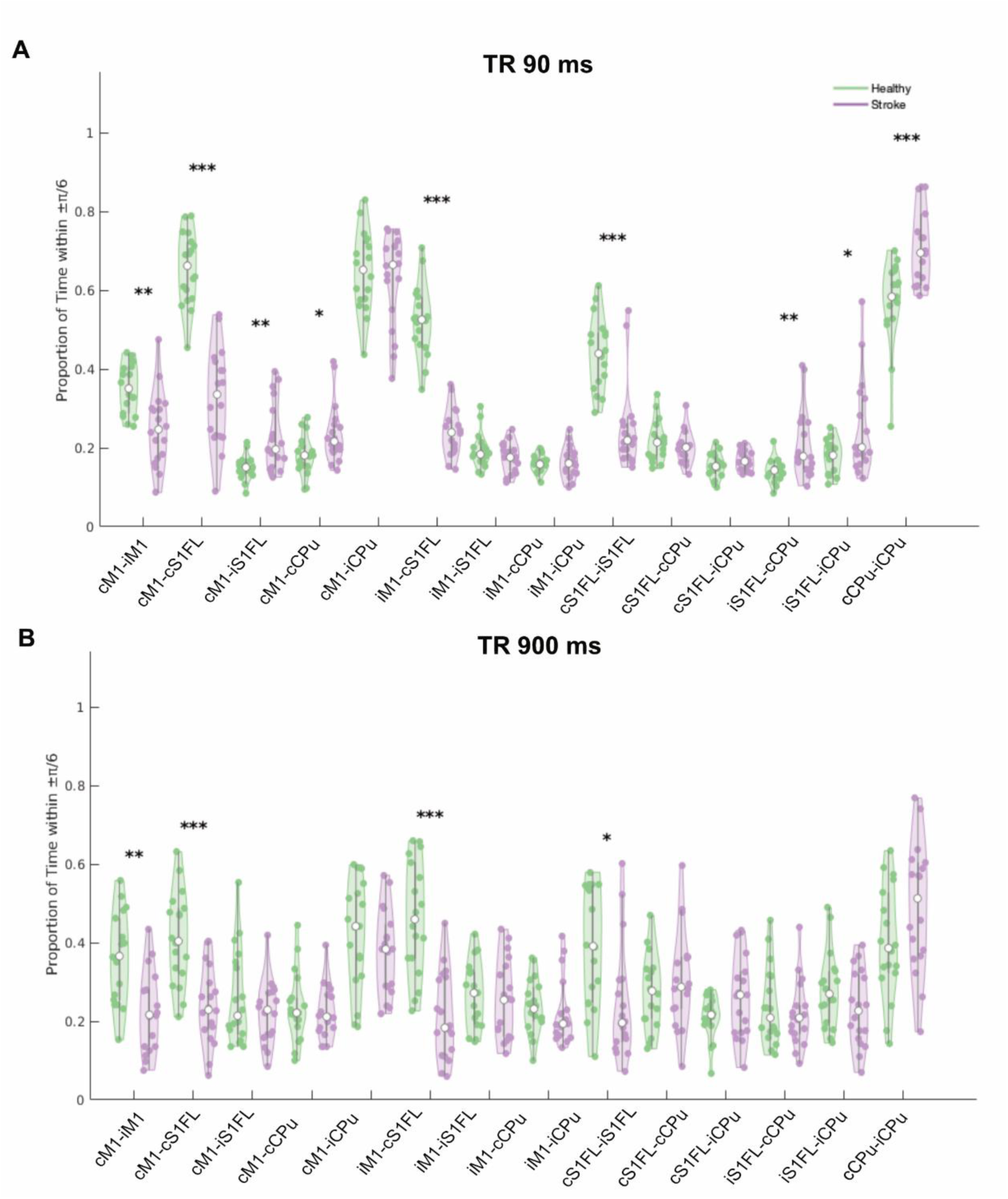
Occupancy analysis. **(A)** Phase occupancy measured as the proportion of time phase differences remained within π/6 across ROIs in ultrafast rs data. Cortical ROI pairs in the healthy group (green) show higher occupancy in a “high synchrony” phase state compared to the stroke group (purple), whereas the subcortical and cortico-subcortical ROI pairs in the stroke group show higher occupancy, compared with the healthy group. **(B)** Phase occupancy measured as the proportion of time phase differences remained within π/6 across ROIs in the downsampled data. No cortico-subcortical or subcortical alterations in occupancy were significant. A two-sample t-test across scans was performed, followed by false discovery rate (FDR) correction for multiple comparisons. (* *P* < 0.05; ** *P* < 0.01; *** *P* < 0.001).

